# The Th1 cell regulatory circuitry is largely conserved between human and mouse

**DOI:** 10.1101/2021.01.11.426266

**Authors:** Stephen Henderson, Venu Pullabhatla, Arnulf Hertweck, Emanuele de Rinaldis, Javier Herrero, Graham M. Lord, Richard G. Jenner

**Affiliations:** Bill Lyons Informatics Centre, UCL Cancer Institute and CRUK UCL Centre, University College London, London, UK; NIHR Biomedical Research Centre at Guy’s and St Thomas’ Hospital and King’s College London, London, UK; School of Immunology and Microbial Sciences, King’s College London, London, UK; Faculty of Biology, Medicine and Health, University of Manchester, Manchester, UK; Regulatory Genomics Group, UCL Cancer Institute and CRUK UCL Centre, University College London, London, UK; Oxford Gene Technology, Oxford, UK; Sanofi, Cambridge, MA, USA

## Abstract

Gene expression programmes controlled by lineage-determining transcription factors are often conserved between species. However, infectious diseases have exerted profound evolutionary pressure, and therefore the genes regulated by immune-specific transcription factors might be expected to exhibit greater divergence due to exposure to species-specific pathogens. T-bet (Tbx21) is the immune-specific lineage-defining transcription factor for T helper type I (Th1) immunity, which is fundamental for the immune response to intracellular pathogens but also underlies inflammatory diseases. We therefore compared T-bet genomic targets between mouse and human CD4^+^ T cells and correlated T-bet binding patterns with species-specific gene expression. Remarkably, we show that the vast majority of T-bet regulated genes are conserved between mouse and human, either via preservation of a binding site or via an alternative binding site associated with transposon-linked insertion. We also identified genes that are specifically targeted by T-bet in humans or mice and which exhibited species-specific expression. These results provide a genome-wide cross-species comparison of T-bet target gene regulation that will enable more accurate translation of genetic targets and therapeutics from pre-clinical models of inflammatory disease into human clinical trials.

## INTRODUCTION

The differentiation of naïve CD4^+^ T cells into T helper type 1 (Th1) effector cells tailors the immune response to target intracellular bacteria and viruses and is critical for effective anti-tumour responses. However, inappropriate Th1 effector cell activation contributes to the development of autoimmune and inflammatory diseases.

The differentiation of Th1 cells from naïve CD4^+^ T cells is controlled by the lineage-determining transcription factor T-bet. Experiments in genetically modified mice have revealed that T-bet is necessary and sufficient for Th1 differentiation [1, 2]. T-bet directly activates Th1 genes such as those encoding the inflammatory cytokines IFNγ and TNF and receptors such as TIM3 (encoded by *HAVCR2*) and CCR5 [3–9]. At these genes, T-bet binds to extended *cis*-regulatory regions (super enhancers) [9, 10] and recruits Mediator and P-TEFb to activate transcription [11]. T-bet also interacts with the H3K4 methyltransferase SETD7 and the H3K27 demethylase KDM6B (JMJD3), recruiting these factors to *Ifng* [12]. Genetic variation at T-bet binding sites is associated with differences in T-bet occupancy between human individuals, including at causal variants associated with inflammatory disease [13], suggesting that differences in T-bet binding between individuals directly contributes to disease risk.

Much of our understanding of Th1 cell function in health and disease comes from studies in mice but the degree to which these findings can be applied to humans is unclear, especially given the evolutionary pressure on the immune system exerted by pathogens [14, 15]. Comparison between the expression profiles of human and mouse T cells over 48 hours of *in vitro* activation with anti-CD3/CD28 has revealed that the T cell activation program is generally shared, but that significant differences do exist between the two species [14, 16]. This variation may be due to differences in transcription factor binding between species. As an example of this, we have previously found that the gene encoding the colon homing receptor GPR15 is only occupied by GATA3 and expressed in human and not mouse Th2 cells [17]. In mouse, expression of the gene is instead specific to Th17 and as peripherally-derived induced regulatory T cells (pTREG), resulting in differences in the types of T helper cells that are trafficked to the human versus mouse colon. Thy-1 (CD90), a GPI-linked Ig superfamily molecule of unknown function is used as a T cell marker in mice but, in humans, it is only expressed on other cell types, potentially depending on the presence or absence of an Ets-1 binding site in the third intron of the gene [18].

However, although differences in transcription factor occupancy have been compared systematically between humans and mice in other cell types, notably hepatocytes [19], such analyses have not been performed for immune cells. Thus, the similarities and differences between human and mouse Th1 cell transcriptional programs remain unknown.

We hypothesised that comparison of T-bet binding sites between human and mouse would determine the degree to which the Th1 transcriptional program is shared between the two species. Here, we present a systematic comparison between T-bet binding sites and gene expression between human and mouse Th1 cells. Surprisingly, given the evolutionary pressure on immune system function, we show that the vast majority of T-bet target genes are shared between the two species. However, we also identify a high-confidence set of genes that are specifically occupied by T-bet in either humans or mice. Species-specific T-bet binding sites are enriched for transposable elements, consistent with a role for these elements in the evolution of immune regulatory sequences between human and mouse. This work defines for the first time both the shared and the divergent aspects of the Th1 transcriptional program between human and mice and provides a framework to support the translation of putative therapeutic pathways identified in murine pre-clinical models into effective treatments for human diseases driven by aberrant Th1 immunity.

## RESULTS

### Identification of shared and species-specific T-bet binding sites

We sought to identify the degree to which T-bet binding sites and target genes were conserved between human and mouse Th1 cells. We first identified the genome positions occupied by T-bet in each species at high confidence (q<0.01 in all replicate ChIP-seq datasets; S1 Table). We then identified the subset of these regions that could be compared between species using liftOver [20]. Conserved binding sites were defined as those present at high confidence in both species and species-specific sites as those present at high confidence in one species and for which there was no evidence of binding in the other species (q>0.1 in all replicates). Binding sites outside of these criteria were judged as indeterminate and were not considered further. This process revealed that around one-third of T-bet binding sites were conserved between species (36% in humans, 32% in mouse) and around two-thirds of sites were species-specific.

To compare T-bet gene targeting between human and mouse, we focused on T-bet binding sites associated with orthologous genes. We found that 2191 genes were occupied by T-bet in either human or mouse. At the majority (1521, 69%) of these genes, a specific T-bet binding site was conserved between species (Conserved, Figs 1A and 1B, S2 Table). These genes included the classical Th1 genes *IFNG, CXCR4, FASLG, HAVCR2, IL12RB2, IL18R1, IL18RAP* and *TNF* (Fig 1C). An additional 349 genes (16%) were also bound by T-bet in both species but at alternative species-specific sites (Alternative, Figs 1A and B, S2 Table), including the genes *TNFSF15, CCR2, MGAT4A* (Fig 1C). Finally, 171 genes (8%) were only bound by T-bet in humans (Hs-specific) and 150 genes (7%) were only bound by T-bet in mouse (Mm-specific; S2 Table). Hs-specific T-bet target genes included *GREM2, TIMD4* and *PKIA*, while Mm-specific T-bet target genes included *Il18, Serpinb5* and *Bend4* (Fig 1C). Thus, we can draw three conclusions from this analysis. Firstly, the majority of T-bet target genes are conserved between mouse and human. Secondly, loss of a binding site at a gene in one species tends to be accompanied by the appearance of an alternative site at the same gene in the other species. Finally, for only a relatively small number of genes is T-bet binding unique to human or mouse.

**Fig 1.**
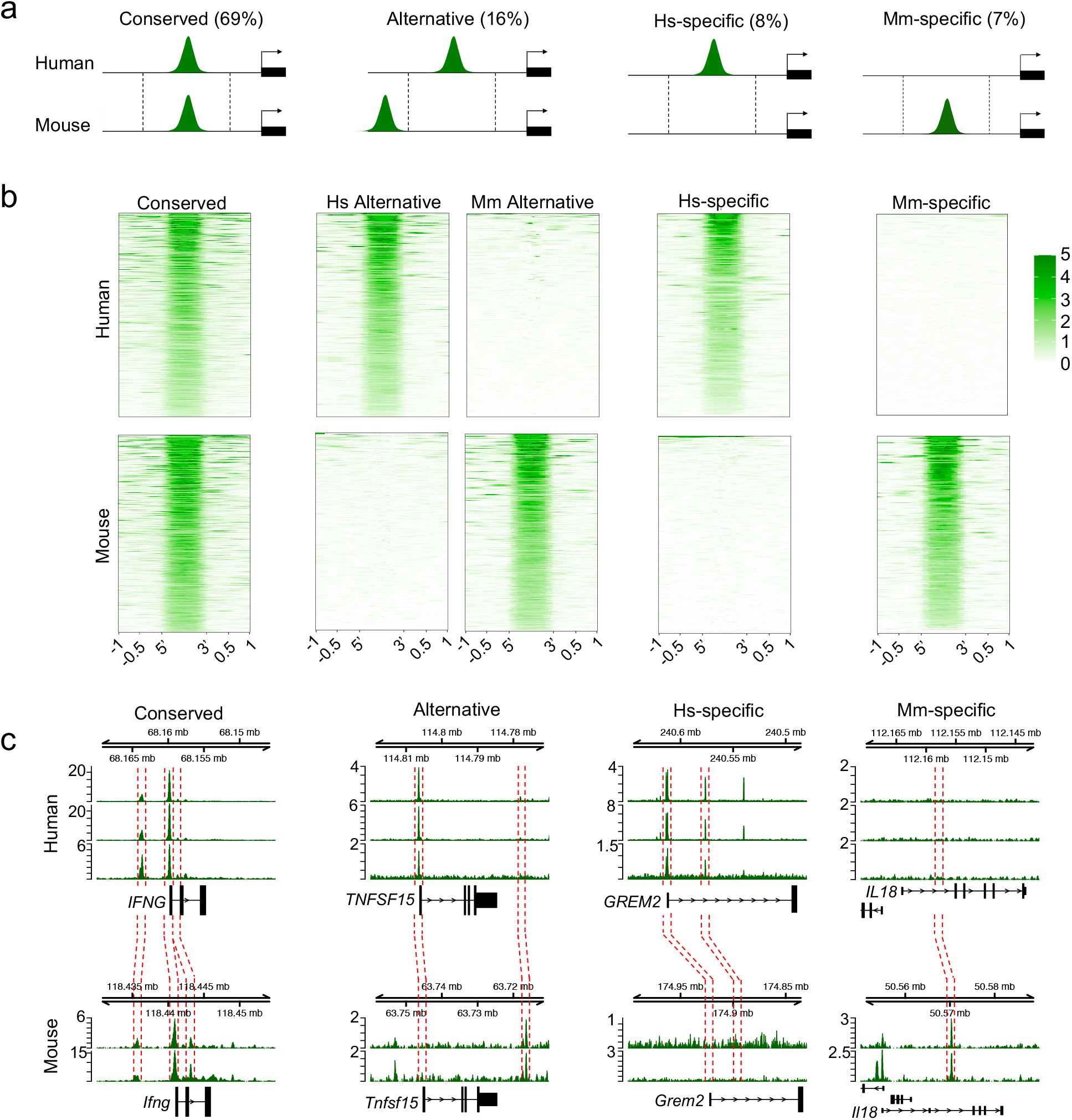
Conserved and specific-specific T-bet binding in human and mouse Th1 cells. **a.** Cartoon showing the 4 different classes of T-bet target genes identified in this study and the proportion of binding sites that fall into each category. Conserved target genes are defined as orthologous genes associated with a high-confidence T-bet binding site at an equivalent location (defined by liftOver) in both species. Alternatively-bound genes are bound by T-bet in both species but at different locations. Hs-specific and Mm-specific target genes are only bound by T-bet in human or mouse, respectively. **b.** Heat maps showing T-bet occupancy at the sets of sites described in a. Sequence reads (per million total reads) at each position are represented by colour, according to the scale on the right. **c.** T-bet binding at example genes with conserved, alternative, Hs-specific and Mm-specific T-bet binding. The red dashed lines show the equivalent locations of T-bet binding sites in the other species, as defined by liftOver.

### Species-specific recruitment of transcriptional co-factors at T-bet binding sites

We next sought to address whether the species-specific T-bet binding sites we identified were likely to be functional. We have previously shown that T-bet recruits P-TEFb, Mediator (MED1 subunit), and the super elongation complex (SEC; AFF4 subunit) to its binding sites in human and mouse Th1 cells [11]. We therefore asked whether species-specific T-bet binding was accompanied by species-specific recruitment of these factors. We gathered ChIP-seq data for these transcriptional regulators in human and mouse Th1 cells and plotted the occupancy of the factors at conserved, alternate and species-specific sites. We found that all of the factors were enriched at conserved sites in human and mouse, consistent with T-bet recruiting these factors in both species (Fig 2 and S1 Fig). In contrast, species-specific T-bet binding sites were only occupied by P-TEFb, AFF4 and MED1 in the species in which T-bet was bound (Fig 2 and S1 Fig). This was also the case for genes bound by T-bet at alternative sites in humans and mouse, with the co-factors only occupying the sites at which T-bet was present in that species. Thus, we conclude that species-specific T-bet binding results in species-specific recruitment of T-bet dependent co-factors.

**Fig 2.**
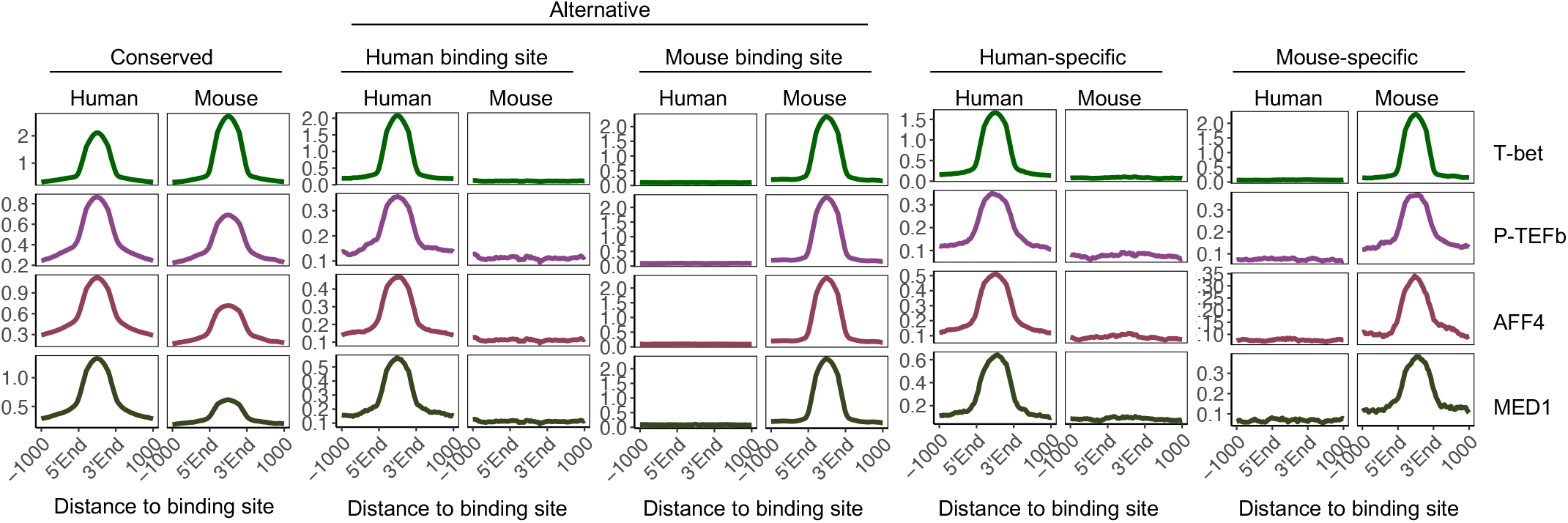
Species-specific T-bet binding is associated with species-specific recruitment of P-TEFb, the super elongation complex and Mediator. Average number of ChIP-seq reads (per million total reads) for T-bet and its co-factors P-TEFb, the super elongation complex subunit AFF4 and the Mediator subunit MED1 across conserved, alternative, human-specific and mouse-specific T-bet binding sites in human and mouse Th1 cells.

### Species-specific T-bet binding is associated with species-specific gene expression

We next sought to determine whether these patterns of T-bet and co-factor binding were associated with differences in gene expression between species. To avoid potential issues with differences in expression being dataset-dependent rather than species-dependent, we performed differential gene expression analysis between human and mouse using 3 independent Th1 cell RNA-seq datasets for each species (Fig 3A, S2 Fig and S3 Table). We found that genes associated with conserved binding sites exhibit similar expression levels in human and mouse (mean log_2_ human/mouse expression ratio of −0.59, std. dev 1.59). Genes bound by T-bet in both species but at alternative sites exhibited more variable expression between species (std dev. 2.46, F 0.42, p<2e^−16^), but a similar mean log_2_ human/mouse expression ratio (−0.21). Thus, loss of T-bet binding during evolution can be functionally neutral as long as the binding site is replaced by an alternative T-bet binding site at the gene. In contrast, genes bound by T-bet specifically in human tended to be more highly expressed in human (mean log_2_ Hs/Mm of 1.95, p=4.4e^−13^, t-test vs Conserved) and, reciprocally, the genes specifically bound by T-bet in mouse tended to be more highly expressed in mouse (mean log_2_ Hs/Mm of −1.55, p=0.0011). Whilst human-specific T-bet target genes constituted 8% of T-bet target genes, they made up 53% (26 of 49) of the T-bet target genes most highly expressed in human vs versus mouse (log_2_ Hs/Mm >5). Similarly, although mouse-specific T-bet target genes constituted 7% of T-bet target genes, they accounted for 63% (22/35) of the T-bet target genes most highly expressed in mouse versus human (X^2^-test, both p<2e^−16^). Thus, species-specific T-bet occupancy is associated with species-specific gene expression. Genes specifically bound by T-bet in humans and significantly (p<1e^−4^) more highly expressed in human versus mouse Th1 cells included *GREM2, TIMD4, TNFSF12* and *PKIA* (Fig 3B). Reciprocally, genes specifically bound by T-bet in mouse and overexpressed in mouse versus human Th1 cells included *Bend4*, *Spata2* and *Serpinb5* (Fig 3B). We conclude that species-specific T-bet occupancy is associated with species-specific gene expression.

**Fig 3.**
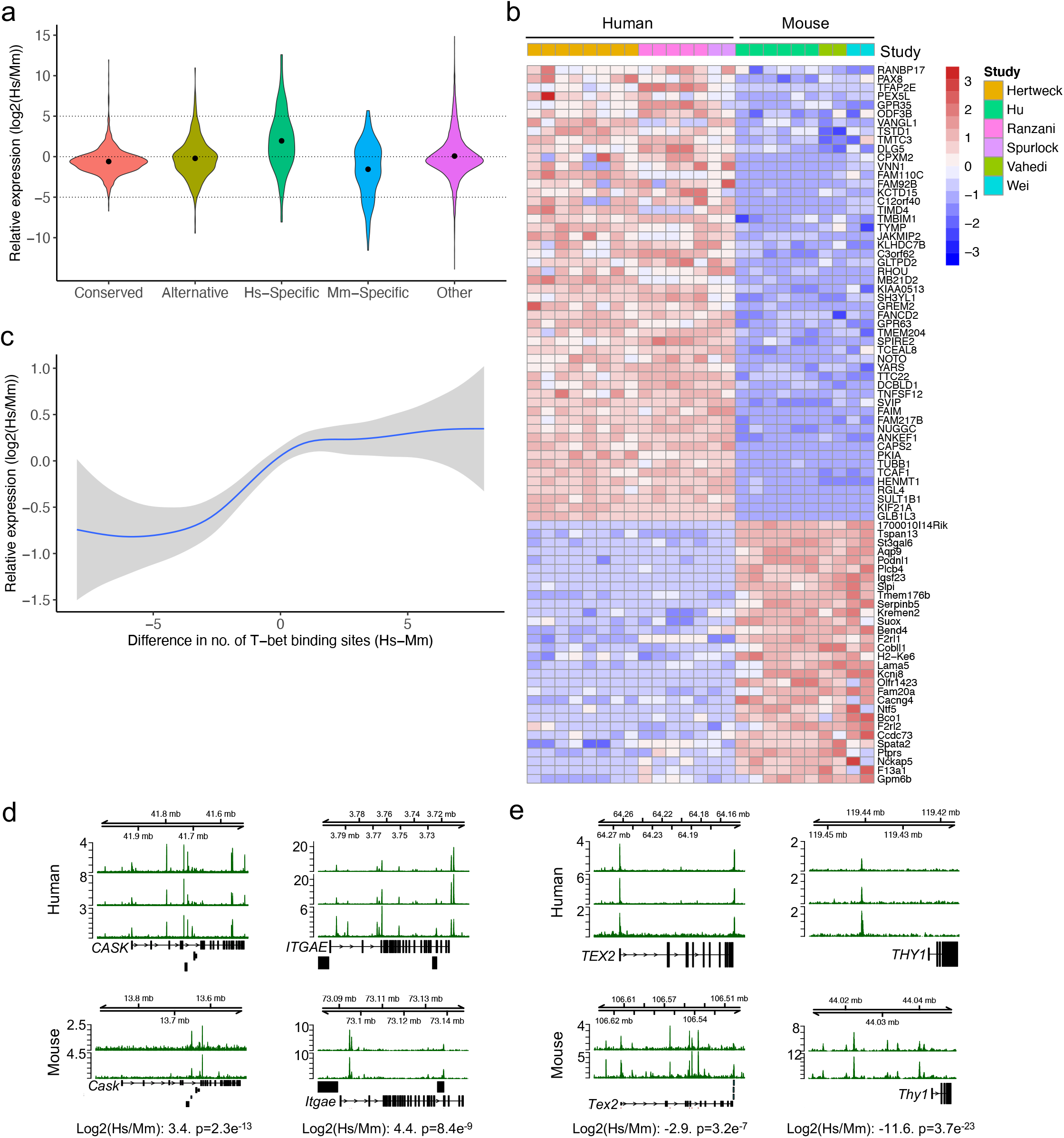
Species-specific T-bet binding is associated with species-specific Th1 gene expression. **a.** Violin plot of the distribution of log_2_ human vs mouse Th1 cell expression ratios for gene sets defined in Fig 1A or at other genes. Median values are marked by a dot. Log_2_ Hs/Mm ratios of 5, discussed in the text, are indicated by dashed lines. **b.** Heatmap showing expression of Hs-specific and Mm-specific genes that are significantly differentially expressed between human and mouse Th1 cells (Welch’s t-test: unadjusted p<1e^−4^). Log_2_ human vs mouse expression ratio is indicated by colour according the scale to the right. The study from which each dataset was taken is indicated by the coloured bar at the top, with the key to the far right. **c.** Loess regression fit of the relation between the log_2_ difference in human and mouse Th1 gene expression and the difference between the number of human and mouse T-bet binding sites for genes bound by T-bet in both species (grey area is the 95% confidence interval). Genes with a greater number of T-bet binding sites in human tend to be more highly expressed in human, and vice versa. **d.** Examples of genes with more T-bet binding sites and which are significantly more highly expressed in human than mouse Th1 cells. **e.** Examples of genes with more T-bet binding sites and which are significantly more highly expressed in mouse than human Th1 cells.

We next considered whether there were any features of T-bet occupancy that could explain why some genes with conserved T-bet binding sites in human and mouse were nevertheless differentially expressed between the species. We found that at these genes, differences in the total number of T-bet binding sites between species was associated with differences in gene expression, with the gene being more highly expressed in the species in which it was bound at the greater number of sites (linear regression of mean log_2_ Hs/Mm expression units per T-bet binding site, p<2e^−6^; Fig 3C). This is consistent with previous observations that Th1 gene expression is driven by T-bet binding to multiple sites across extended *cis*-regulatory regions, later termed super-enhancers [9–11]. Genes bound by T-bet at a greater number of sites and more highly expressed in humans than in mouse included *CASK*, *ITGAE* and *GZMK* (Fig 3D and S3 Table), while genes bound by T-bet at a greater number of sites and more highly expressed in mouse included *Thy1* (consistent with its expression in mouse but not human T cells), *Tex2* and *Nfatc1* (Fig 3E and S3 Table). Thus, in addition to the absolute presence and absence of T-bet, the relative number of T-bet binding sites is also associated with differential expression of Th1 genes between species.

### Species-specific T-bet binding correlates with the presence or absence of a T-bet DNA binding motif

Like other T-box transcription factors, T-bet binds a specific DNA sequence motif [8, 9]. We therefore considered whether differences in T-bet binding between human and mouse might be related to differences in the sequences at those sites between the species. To address this, we first identified consensus motifs enriched in the complete sets of high-confidence T-bet binding sites in human and mouse. This confirmed enrichment of highly similar motifs that matched the previously determined T-bet DNA binding motif [8]) in both species (Hs p=1e^−623^; Mm p=1e^−642^; Fig 4A).

**Fig 4.**
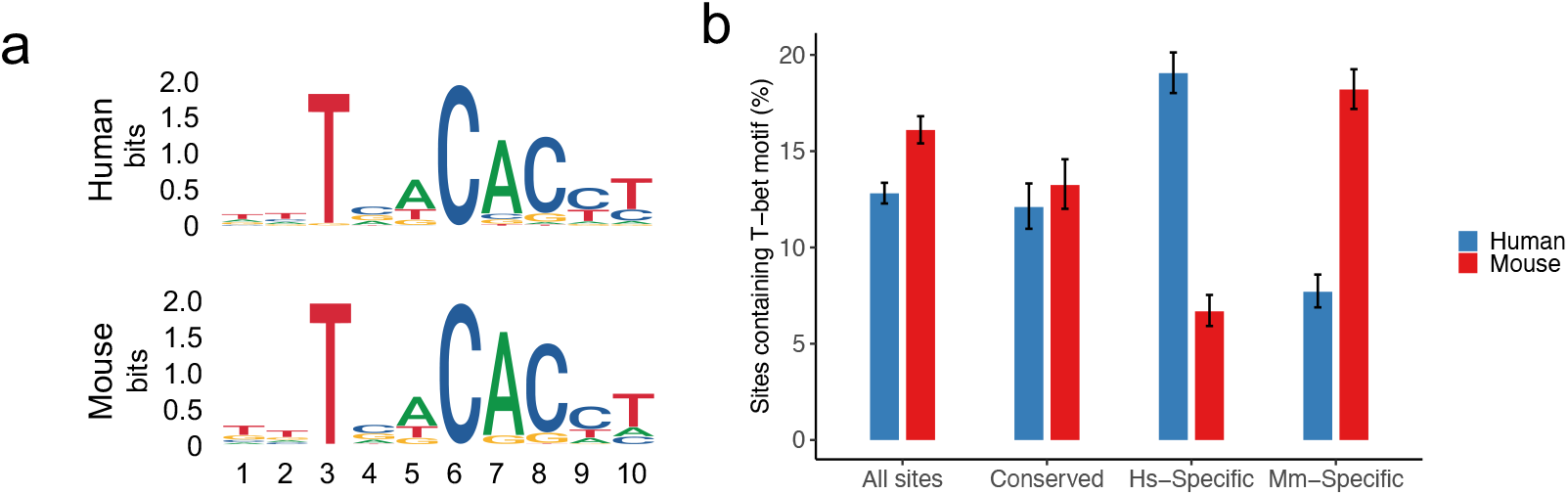
Species-specific T-bet binding correlates with the presence or absence of the consensus T-bet DNA binding motif. **a.** DNA binding motifs matching a previously identified consensus T-bet DNA binding motif [8] enriched in the set of T-bet binding sites in human (top) and mouse (bottom) Th1 cells.**b.** Proportion of all T-bet binding sites, conserved T-bet binding sites, Hs-specific T-bet binding sites and Mm-specific T-bet binding sites in human and mouse that contain a sequence matching the consensus T-bet DNA binding sequence in that species shown in a (error bars represent the 95% confidence interval of the binomial test).

We then used FIMO [21] to quantify the proportion of conserved and species-specific T-bet binding sites that contained the T-bet DNA binding motif. We found that the motif could be identified with confidence at roughly equal proportions of conserved T-bet binding sites in human and mouse (12.1% and 13.2%, respectively; Fig 4B). In contrast, 19.1% of human-specific T-bet binding sites contained a T-bet binding motif in human and this dropped to 6.7% for the equivalent loci in mouse (Fig 4B). Reciprocally, 18.2% of mouse-specific T-bet binding sites contained a T-bet binding motif in mouse and this dropped to 7.7% in human. Thus, whether or not T-bet binds to a genomic location in human versus mouse correlates with whether or not the T-bet DNA binding motif is present, suggesting that many of the differences in T-bet binding between species is due to sequence divergence at these sites.

### Transposable elements are enriched at species-specific T-bet binding sites

Transcription factor binding sites can be located within transposable elements (TEs) and TE invasions have been postulated to contribute to the evolution of regulatory gene networks [22]. We therefore considered that TEs may have played a role in the diversification of T-bet binding sites between human and mouse. To test this, we compared the proportions of the different categories of T-bet binding sites that overlapped TEs of both species (Fig 5A). First looking at conserved binding sites, we found that only 3% overlapped a TE in humans and <1% in mouse. In comparison, 10% of human-specific and 5% of mouse-specific binding sites overlapped a TE. Similarly, 10% of alternative sites bound by T-bet in human and 5% of alternative sites bound by T-bet in mouse overlapped a TE. The enrichment of TEs at species-specific binding sites were highly significant both with a chi-square test (Hs *X*^2^ = 67.5, Mm *X*^2^ = 60.5, both p<2e^−14^) and with permutation tests (n=10,000, p<1e^−5^) (Fig 5B). The association of alternative and species-specific binding sites with TEs was not an artefact of the genomic distribution of these sites; although species-specific T-bet binding sites were enriched at distal locations compared to other T-bet binding sites (Kruskal-Wallis test, Hs p<1e^−4^, Mm p<1e^−3^,) TEs were not enriched at distal sites (p=0.07; S3 Fig). Breaking down TEs into their different classes revealed some differences between human and mouse, with LINE1s and LTRs being associated with both human and mouse-specific binding sites, while LINE2s were more strongly associated with alternative sites in humans and SINEs more strongly associated with alternative sites in mouse (Fig 5B). Thus, these data are consistent with TE activity contributing to the divergence of T-bet binding sites between human and mouse.

**Fig 5.**
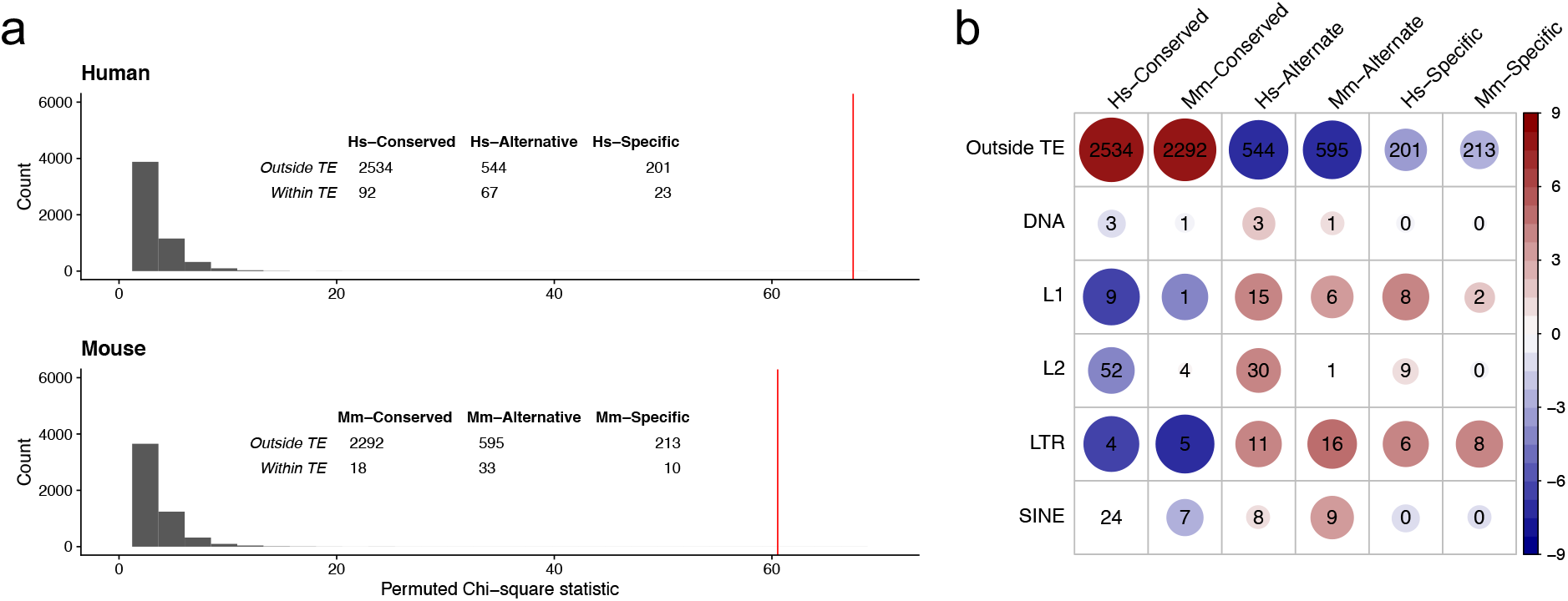
Species-specific T-bet binding sites are associated with transposable elements. **a.** Permutation test of the association between binding site types and TEs. In both human and mouse, Alternative and species-specific binding sites are more likely to overlap a TE than Conserved binding sites. The red bars show the observed overall *X*^*2*^ of the inset table. The histogram shows a *X*^*2*^ null-distribution based on 10,000 permutations of the data. **b.** Heatplot of the chi-square overlaps of the different classes of T-bet binding sites with different classes of TEs. The numbers show the raw table data, colour represents the standardised residuals of the *X*^*2*^ table data, according to the scale on the right, and circle size represents the absolute standardised residual value.

## DISCUSSION

We have determined the degree to which the Th1 cell regulatory circuitry is conserved between human and mouse. We have found that the majority of T-bet binding sites are conserved between species and that T-bet target genes associated with conserved binding sites tend to exhibit similar levels of expression. At genes with conserved binding sites, the presence of additional T-bet binding sites in human or mouse is associated with increased expression in that species. For genes at which T-bet binding sites are not conserved, it is most often the case that an alternative binding site is present at a different position in the other species and gene expression is maintained. At only a minority of genes is T-bet binding unique to human or mouse and these genes tend to be more highly expressed in the species in which T-bet is bound.

Species-specific binding sites overlap TEs, suggesting that transposition of these elements has played a role in the divergence of the Th1 cell regulatory circuitry between human and mouse.

Our analysis was designed to minimise the number of false-positive binding sites. We only considered binding sites identified at high-confidence (q<0.01 in all replicates) and only judged a site to be species-specific if the region could be identified in the other species (by liftOver) and there was absolutely no evidence of binding (q>0.1 in all replicates). Application of these criteria revealed that around one-third of human and mouse T-bet binding sites were conserved in the other species. The degree to which transcription factor binding is conserved between human and mouse immune cells has not previously been determined but similar data are available for embryonic stem cells, hepatocytes and liver and various cell lines. Comparison between the degree of conservation of binding sites between T-bet and those of other transcription factors is not straightforward because of differences in the criteria used to assign a position as bound or not bound between studies. However, the degree of conservation we found for T-bet is similar to that previously found for master regulator HNF transcription factors in hepatocytes [19]. Given that the immune system is subject to continuous evolutionary pressure in the form of rapidly evolving pathogens [23], the similar levels of binding site conservation between T-bet and HNF transcription factors is perhaps unexpected but reinforces the notion that the regulatory circuitry underlying the specification of T helper cell lineages is highly conserved between species.

Although one-third of T-bet binding sites are conserved between human and mouse, the proportion of T-bet target genes that are conserved is higher, with 85% of T-bet target genes bound in both species. For the vast majority of these genes, the location of at least one T-bet binding site was conserved. Furthermore, for the majority of genes at which a T-bet binding is lost during evolution, an alternative binding site arises at the same gene. This suggests considerable pressure to conserve T-bet binding sites during human and mouse evolution.

Divergence in T-bet binding between species is correlated with divergence in gene expression. Genes specifically bound by T-bet in human or mouse exhibit higher expression in the species in which the gene is bound; genes bound by T-bet only in humans tend to be expressed more strongly in humans and vice versa. Differences in the number of T-bet binding also correlates with differential expression of T-bet target genes that are shared between species, with the acquisition of additional T-bet binding sites associated with increased expression of the gene in that species. This suggests that the number of T-bet binding sites at a gene has been subjected to selective pressures and is consistent with evidence showing that transcription factor binding sites can regulate gene expression in an additive fashion [9, 10, 24].

Whether or not T-bet was detected at sites in human and mouse was correlated with the presence or absence of a T-bet binding motif. This indicates that DNA sequence mutations underlie some of the divergence in T-bet binding between human and mouse. However, the association between T-bet binding and presence of the binding motif was not complete, with some binding sites lacking the consensus T-bet binding sequence, while other sites contained the binding sequence but lacked T-bet occupancy. This indicates that other variables, for example motifs for co-binding transcription factors or differences in chromatin state, may also contribute to variation in T-bet binding between species.

We found that species-specific T-bet binding sites were enriched for association with TEs, especially LINE1 and LTRs. Alternative binding sites were also associated with these elements and additionally with LINE2 in human and SINEs in mice. TEs have previously been reported to be co-opted as regulatory elements by their host and to contain bindings sites for transcription factors [22]. TEs have also been found to be associated with species-specific binding of transcription factors including STAT1/IRF1, TP53 and OCT4/NANOG [25–27], and to the establishment of enhancers at innate immune genes [27]. Thus, in discovering enrichment of TEs at T-bet binding sites, our study expands the contribution of TEs to adaptive immune cell regulatory programs.

In summary, by comparing T-bet binding and gene expression between human and mouse, we have found that the Th1 regulatory circuitry is generally conserved between species but that some key differences exist. These data will be of value in guiding the appropriate use of mice for target identification and drug development for human inflammatory diseases.

## METHODS

### Comparison of T-bet binding data between human and mouse

Human and mouse Th1 cell ChIP sequencing data was downloaded from GEO (Hs T-bet rep1=GSM2176976, Hs T-bet rep2=GSM2176974, Hs T-bet rep3=GSM776557, Mm T-bet rep1=GSM998272, Mm T-bet rep2=GSM836124, Hs P-TEFb=GSM1527693, Mm P-TEFb=GSM1527702, Hs AFF4=GSM1961563, Mm AFF4=GSM1961559, Hs MED1=GSM1961567, Mm MED1=GSM1961557). After trimming low quality reads from both ends using seqtk (error rate threshold 0.05), reads were aligned to the GRCh38 or GRCm38 assemblies using Bowtie2 with default “sensitive” settings (-D 15 -R 2 -N 0 -L 22 -i S,1,1.15). High-confidence T-bet binding sites were identified by comparison to input using MACS2 (q<0.01). A high confidence set of binding sites for each species was then defined as the binding site coordinates that overlapped in all replicates. Similarly, low-confidence T-bet binding sites for each species were defined as those identified by MACS2 at q<0.1 in any replicate. Binding sites that overlapped ENCODE blacklist regions (https://github.com/Boyle-Lab/Blacklist/tree/master/lists) were removed. The coordinates of high-confidence binding sites were extended by 1 kb either side and the equivalent coordinates identified in the other species using the mm10tohg38 and hg38tomm10 liftOver chains (from UCSC) and the *rtracklayer* package for R. Equivalent location was defined as a single range from the beginning to the end of the lift-over. Conserved binding sites were defined as those present at high confidence in both species and species-specific sites as those present at high confidence in one species and for which there was no evidence of binding in the other species (q>0.1 in any replicate).

T-bet binding sites were associated with the nearest gene as defined by human GENCODE V29 transcripts or mouse GENCODE M20 transcripts annotations. These transcript models were chosen for compatibility with the GRCh38 or GRCm38 assembly, the Ensembl gene ortholog models, and available UCSC genome browser tracks. Orthologous genes were identified using Ensembl Compara and downloaded via Ensembl Biomart. Genes with conserved T-bet binding sites were defined as those associated with conserved sites in both species. Genes with alternate binding sites were defined as those associated with species-specific binding sites in both species and no conserved sites. Genes with species-specific binding were defined as those associated with a high-confidence T-bet binding site in one species and no binding sites in the other species.

### Visualisation of ChIP-seq data

We used ngsplot [28] to extract read coverage around binding sites and the equivalent regions in the other species from a single merged ChIP BAM alignment file for each species and to generate average binding profiles (metagenes) and heatmaps (both showing read counts per million mapped reads). To visualise T-bet binding data at individual genes, we used deeptools bamcoverage [29] to create bigwig files (read counts per million mapped reads) (CPM) and then plotted these in their genomic context using the Gviz tool for R [30].

### Gene Expression

RNA-seq data were downloaded from GEO in fastq format; human Th1 datasets from [31, 32ÕHertweck, 2016 #1078], mouse Th1 datasets from [33–35]. Gene centred expression estimates were made using *kallisto* [36] together with the GENCODE V29 (human) and M20 (mouse) transcript models. Human and mouse expression estimates were then modelled separately using *DESeq2* [37], with experimental source and cell type (Th1/Th2) treated as covariates for batch correction (RNA type (polyA+ vs total RNA) was also tested but found to be a negligible effect). Once normalised and batch corrected, human and mouse expression data were similar in distribution, but prior to cross species comparison the data was zero centered. Expression heatmaps were drawn with variance stabilising transformations (*vst*) of the data for a more easily interpretable colour scale. Linear regression was used to calculate the significance of the association between the difference in the number of T-bet binding sites between species (−5 to +5) and the log_2_ human vs mouse expression ratio.

### Motif Analysis

A consensus motif matching the previously identified T-bet DNA binding motif [8] was identified in the complete set of high-confidence human and mouse T-bet binding sites with the findMotifsGenome.pl programme from the HOMER tools suite [38] using the following parameters: hg38 or mm10 –size given –mask. Conserved and species-specific T-bet binding sites were identified as before, except without extending regions by 1 kb before liftOver. The human or mouse position-weight matrix was then used to identify the T-bet consensus motif within the different sets of binding sites (all, conserved, species-specific) for that species the using the FIMO tool from the MEME suite [21]. Confidence intervals were calculated using prop.test in R.

### Transposable elements

Coordinates of human and mouse transposable elements were downloaded from the UCSC hg38 and mm10 Table Browser. Specifically, we used the nested repeat tracks from Repbase [39], which merge closely adjacent fragmented or nested repeats into single elements. Binding sites were defined as overlapping a TE if the central 40 bp region was fully enclosed by a TE. Tests of independence were carried out using the R chisq.test function (*X*^*2*^). As the numbers of Conserved, Alternate and Specific sets of binding sites were quite different, we used a permutation test to double-check the observed *X*^*2*^ value by comparing it to 10,000 permutations of the TE labels. To represent more clearly the complex associations between gene sets and particular TE types (e.g SINE, LTR, etc), we plotted the standardised residuals of their *X*^*2*^ test of independence table (Observed – Expected / √ Expected); the standardised metric is cognisant of the wide difference in size of the gene sets. Distances between binding sites or TEs and the nearest gene were taken from the mid-point of the binding sites or TE to the gene TSS.

## ACKNOWLEDGEMENTS

This work was funded by MRC grants to RGJ and GL (MR/M003493/1 and MR/R001413/1)), the Cancer Research UK-University College London (CRUK-UCL) Centre (award C416/A25145) and the Guy’s and St Thomas’ Biomedical Research Centre.

## AUTHOR CONTRIBUTIONS

Study conception: RGJ and GML. Experimental design: SH, VP, JH, RGJ. Data analysis: SH, VP, AH, RGJ. Study supervision: EdR, JH, RGJ, GML. Writing of the paper: RGJ, with input from SH, AH, JH and GML.

## SUPPORTING INFORMATION

### Supplemental Figure Legends

**S1 Figure.**
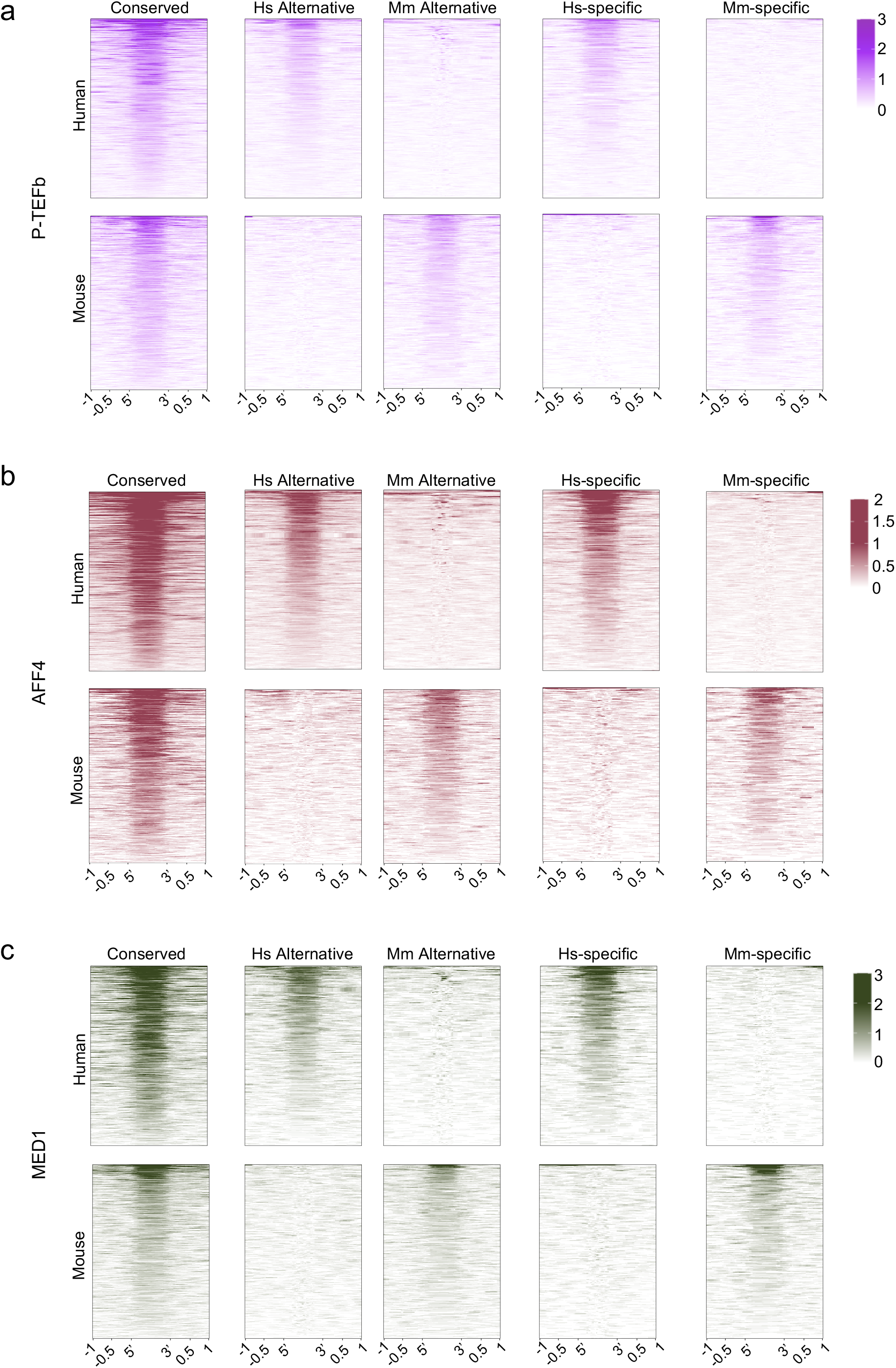
Occupancy of transcriptional co-activators at T-bet binding sites. Heat maps showing P-TEFb, AFF4 and MED1 occupancy at the sets of sites at which T-bet binding is shown in Fig 1B. For each factor, sequence reads (per million total reads) at each position are represented by colour, according to the scales on the right.

**S2 Figure.**
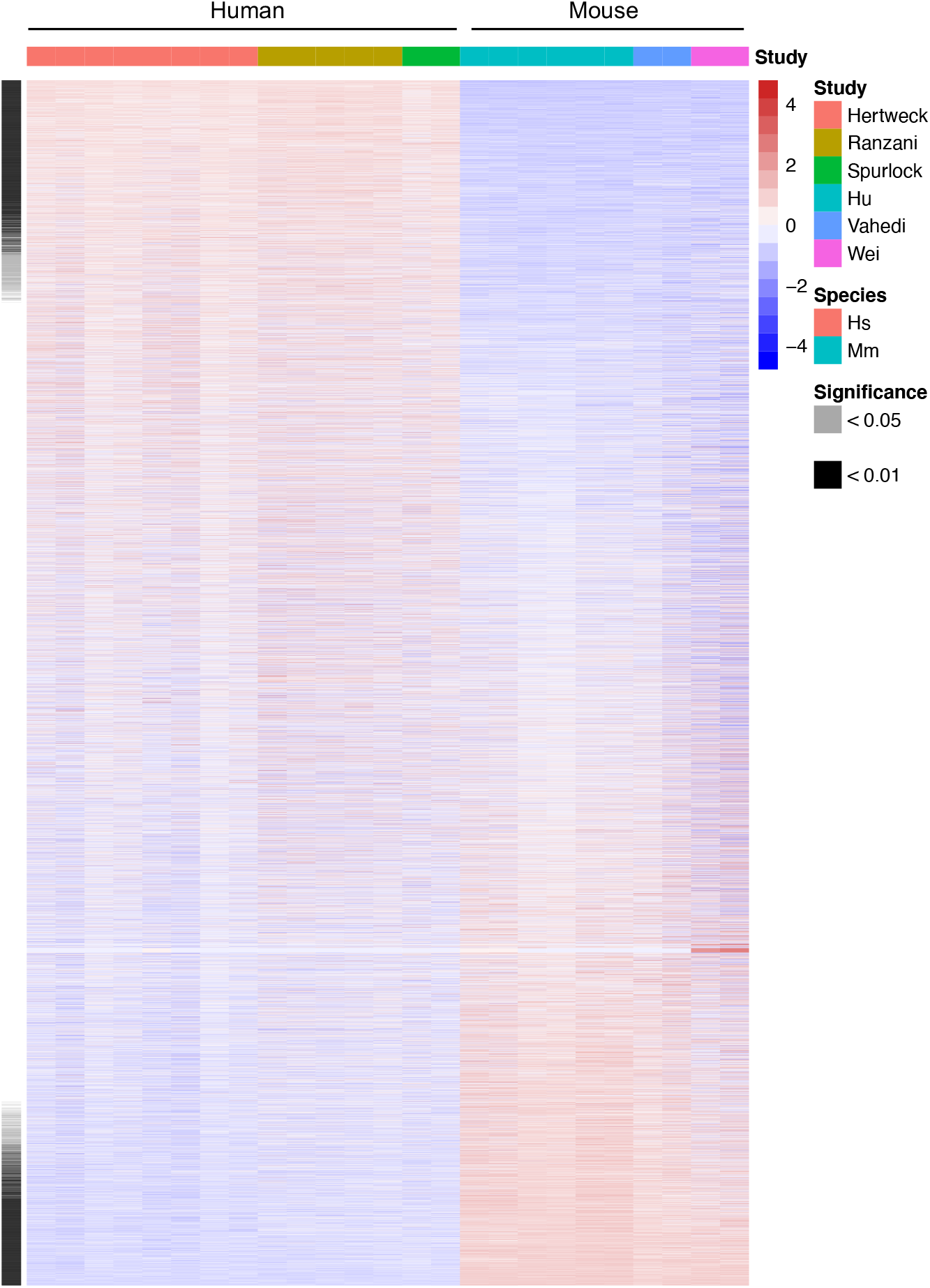
Differential gene expression between human and mouse Th1 cells. Relative expression of orthologous genes between human and mouse Th1 cells. Genes differentially expressed between species are marked (Welch’s t-test, unadjusted p<0.01 or <0.05).

**S3 Figure.**
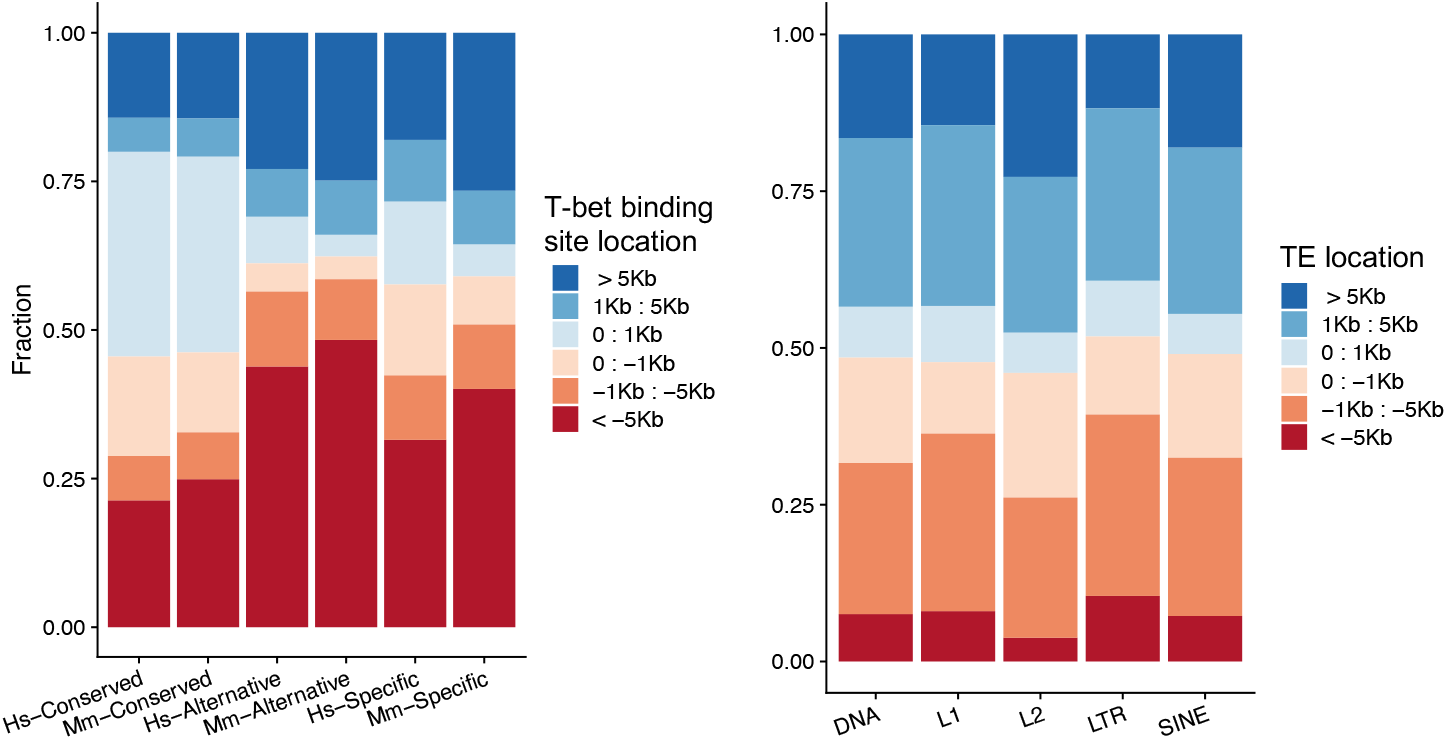
The overlap between species-specific T-bet binding sites and TEs is not merely due to similar genomic distributions. **a.** Stacked bar plots showing the proportion of the different classes T-bet binding sites located at varying distances from the closest gene TSS (− upstream, + downstream). **b.** Stacked bar plots showing the proportion of different classes of TEs located at varying distances from the closest gene TSS (− upstream, + downstream).

### Supplemental Tables

**S1 Table. Human and mouse T-bet binding sites.**

**S2 Table. Conserved, Alternative and species-specific T-bet binding.**

**S3 Table. Human versus mouse Th1 gene expression.**

